# Transcriptional Markers of Organic Substrate Utilization in a Marine Bacterium

**DOI:** 10.64898/2026.06.26.734754

**Authors:** Samantha Cerda, Melanie Cohn, Liang Zhao, Scott Gifford

## Abstract

Marine dissolved organic carbon is a chemically complex substrate pool that fuels heterotrophic bacteria, yet it remains difficult to determine which compounds are used by specific microbes. Bacterial transcriptomes offer a potential biosensor of substrate availability, but the reliability of this approach in chemically mixed substrates remains uncertain. Here, we evaluated the reliability of this transcriptional sensor approach using the model marine bacterium *Ruegeria pomeroyi* DSS-3 grown on either glucose or a defined mixture containing glycerol, benzoate, succinate, leucine, dimethylsulfoniopropionate, and trimethylamine N-oxide. Genome-wide transcription differed strongly between treatments, with the mixed-substrate treatment enriched in genes associated with C1 metabolism, sulfur oxidation, benzoate degradation, and motility. Across substrates, the most diagnostic transcriptional responses occurred at pathway entry points and first committed reactions, including glucose transport and Entner-Doudoroff metabolism, trimethylamine N-oxide transport and catabolism, and early steps of aerobic benzoate oxidation. In contrast, downstream metabolic genes were less substrate-specific, likely because multiple pathways converged on shared central metabolic intermediates. Transporter transcription was also less consistently diagnostic than expected, although substrate-binding subunits often showed the strongest responses within transporter complexes. Comparisons with previous single-substrate studies indicated that some transcriptional markers, particularly benzoate oxidation genes, remained detectable in the substrate mixture, whereas glycerol and succinate responses were weakened or lost. These findings show that transcriptomics can provide useful insight into bacterial substrate use, but interpretation is most robust when focused on experimentally validated transporters and early pathway genes, and when evaluated in the context of pathway connectivity, cellular physiology, and substrate mixture complexity.

**IMPORTANCE:** Microbes help control how carbon moves through the ocean, but it is often difficult to determine which organic compounds individual bacteria are using. This study tested whether gene expression in the marine bacterium *Ruegeria pomeroyi* can be used to identify the types of carbon compounds available in its environment. We found that some genes, especially those involved in initial steps of substrate use, provided clear signals of which compounds were present. However, these signals became harder to interpret when the bacterium was exposed to a mixture of compounds rather than a single substrate. We also found that some gene responses reflected broader physiological or behavioral changes rather than direct use of a specific compound. These results show both the promise and limits of using bacterial gene expression as a biosensor of marine carbon chemistry, and they help identify the kinds of genes most useful for interpreting complex environmental transcriptomic data.

## INTRODUCTION

Bacteria are major processors of dissolved organic carbon in aquatic environments, yet it remains difficult to determine which organic substrates individual taxa use and which genes provide the clearest signatures of that activity. This challenge is especially important in the ocean, where microbial transformations of organic matter regulate carbon cycling, nutrient regeneration, and food web dynamics. Marine dissolved organic carbon is the third largest reservoir of organic carbon on Earth, and by cycling large amounts of carbon while rapidly exchanging it among the atmosphere, biosphere, and ocean interior, the ocean helps sustain ecosystems and regulate the global carbon cycle (DeVries 2022). Despite this importance, the chemical composition, biological availability, and microbial turnover of marine dissolved organic carbon remain poorly resolved.

Multiple sources contribute to the marine dissolved organic carbon pool, including riverine and estuarine inputs, as well as biologically derived compounds released through sloppy feeding, cell lysis, and phytoplankton exudation (Moran et al. 2022). This pool is chemically diverse and analytically challenging to characterize (Catalá, Shorte, and Dittmar 2021; Dittmar and Stubbins 2014). Many labile compounds are also rapidly consumed, allowing them to fuel microbial metabolism while remaining at concentrations that are difficult to detect directly (Bauer, Williams, and Druffel 1992; Jiao et al. 2010; Kirchman et al. 2001; Moran et al. 2022; Raven and Falkowski 1999). Consequently, chemical insight into marine dissolved organic carbon often depends on labor-intensive, low-throughput approaches, some of which recover only a small fraction of the organic molecules present (Chow et al. 2022; Moran et al. 2016).

This chemically diverse dissolved organic carbon pool is transformed by microbial communities whose composition and metabolic activity vary across space, time, and environmental context. Heterotrophic bacteria are central to these transformations, mediating nutrient regeneration, carbon remineralization, and marine food web dynamics (Azam et al. 1983). The flux of labile dissolved organic carbon through heterotrophic bacterioplankton is among the largest organic carbon fluxes in the biosphere (Hansell 2013; Nowinski and Moran 2021), yet it remains difficult to determine which microbes consume which compounds under natural conditions. Approaches that can identify available carbon substrates while simultaneously linking them to microbial activity would therefore provide an important advance for understanding marine carbon cycling.

A transcriptomics perspective offers one route to infer dissolved organic carbon availability from microbial responses. Whereas genomic data describes the organisms present and their metabolic potential, transcriptomic data captures genes being actively expressed under current conditions. Because bacterial messenger RNA inventories are small and turn over rapidly, transcription can provide a near real-time readout of cellular responses to substrate availability (Moran et al. 2013). Transcriptomes may therefore serve as biosensors of the substrates available to microbial communities. However, the extent to which transcriptomic patterns faithfully report substrate availability remains unclear, particularly in chemically complex substrate mixtures.

The heterotrophic marine bacterium *Ruegeria pomeroyi* DSS-3 is a useful model for testing whether transcriptomes can serve as biosensors of substrate availability. DSS-3 is a member of the Roseobacter clade, a metabolically versatile group that is abundant in coastal and open-ocean surface waters and can comprise a substantial fraction of marine bacterioplankton communities (González et al. 2003). This abundance, combined with broad metabolic capabilities, suggests that roseobacters play important roles in marine biogeochemical cycling. DSS-3 is also well suited for this question because its growth physiology, morphology, and genome have been extensively characterized (Rivers et al., 2014). Previous studies have shown that DSS-3 responds transcriptionally to environmentally relevant organic matter inputs. For example, invasion experiments with filtered seawater from seasonal phytoplankton blooms revealed shifts in gene expression involved in carbon, nitrogen, and sulfur metabolism, metal and vitamin metabolism, biotic interactions, and stress responses (Nowinski and Moran 2021). Similarly, prior work with defined media and single added carbon substrates supported the idea that transcriptomic profiles can indicate substrate availability (Boulton 2021). However, much less is known about how these transcriptional signals change when cells are exposed to mixtures of substrates, a more realistic scenario given the enormous chemical diversity of marine dissolved organic carbon (Dittmar 2014).

In this study, we asked whether the transcriptomic response of DSS-3 is a reliable indicator of carbon substrate availability in a mixed-substrate environment and, if so, which genes provide the clearest signals of substrate use. To address this, we cultured DSS-3 in a defined carbon mixture containing glycerol, benzoate, succinate, leucine, dimethylsulfoniopropionate (DMSP), and trimethylamine *N*-oxide (TMAO). This carbon mixture treatment represents several major classes of marine organic substrates, including amino acids, aromatic compounds, and sulfur- and nitrogen-containing methylated molecules. As a control, we grew DSS-3 on glucose alone, a condition expected to emphasize glycolysis and central carbon metabolism rather than the C1-focused metabolism predicted in the carbon mixture. This design allowed direct comparison of transcriptional responses between a single-substrate treatment and a chemically diverse substrate environment in which enriched genes could be linked back to known available compounds. We hypothesized that transporters and genes encoding the first committed steps of metabolic pathways would provide the strongest indicators of substrate availability. We further hypothesized that transcriptional signals previously observed in single-substrate experiments would become less distinct in the carbon mixture but would still retain enough specificity to identify the actively used substrates.

## METHODS

### Carbon substrates solutions

Carbon substrate stock solutions were prepared in Milli-Q water and filter sterilized through 0.2-µm polyethersulfone (PES) filters into sterile, Milli-Q-rinsed Falcon tubes. Filter-sterilized stock solutions were then added to experimental flasks to achieve a final concentration of 6 mM carbon equivalents. For the glucose treatment, D-glucose was added to a final concentration of 1 mM, equivalent to 6 mM carbon. For the carbon-mixture treatment, individual substrates were added to final concentrations of 0.33 mM trimethylamine N-oxide (TMAO), 0.20 mM dimethylsulfoniopropionate (DMSP), 0.14 mM benzoate, 0.25 mM succinate, 0.167 mM leucine, and 0.33 mM glycerol. Each compound contributed approximately 1 mM carbon equivalent, yielding a total of 6 mM carbon equivalents in the mixture.

### DSS-3 cultivation and experimental setup

DSS-3 was streaked from cryostocks onto ½ yeast-tryptone-sea salts (YTSS) agar plates that where incubated in the dark at 29°C. After one day of incubation, one colony from each plate was transferred into 3 mL ½ YTSS liquid media in duplicate (with 2 non-inoculated controls), and inoculated in a mixer in the dark at 29°C, 208 RPM. After 1 day of incubation, 1 mL from each culture was transferred to a 1.7 mL microcentrifuge tube (VWR) and centrifuged for 20 seconds. The supernatant was removed and 1 mL of autoclaved 30 psu NaCl was added and mixed with the cell pellet. This step was repeated twice and the “washed” cell pellet was resuspended with 1 mL of defined-salt minimal basal media (dsMBM, 30 psu) with no organic carbon sources added. The cells from this resuspension were inoculated at a target concentration of 50,000 cells mL^-1^ into sterilized 200 mL Erlenmeyer flasks containing 150 mL of dsMBM liquid media with either the glucose or carbon mixture as the carbon source as described above. The cultures were incubated in the dark at 29°C, 208 RPM.

### Cell Enumeration

Culture growth was monitored by flow cytometry (Guava easyCyte HT, Cytek Biosciences). Samples were diluted as needed with a Tris HCl buffer and stained in the dark for 30 minutes with SYBR Green Nucleic Acid Stain at a final concentration of 20x and ran at a maximum of 30,000 events for 60 seconds. A manually defined gate count was used for cell quantification, and the threshold for Grn-B was 25. Compensation controls are as follows: FSC- 11.8, SSC- 26.9, GRN-B- 21.7, YEL-B- 3.36, RED-B- 6.17. Cell counts for all treatments were subtracted against a SYBR-stained Tris EDTA control.

### RNA collection

When the C-mixture treatment reached stationary phase and the glucose control reached late exponential phase, 30-40 mL of culture from each flask was added to a sterile 50 mL syringe, and gently passed through a 0.22 µm pore, 25 mm PES filter (Fischer, prerinsed with Milli-Q water). After filtering, the filter was placed in a 2 mL cryotube (VWR) and flash frozen in liquid nitrogen for at least 5 minutes before being transferred to -80°C for storage. Total time from cell collection to flash freezing averaged about 4 minutes.

### RNA extraction

RNA was initially extracted using the RNeasy Mini Kit (Qiagen) following the manufacturer’s instructions. Because this approach did not consistently produce the target RNA yields, only the Glucose A sample was extracted with the RNeasy Mini Kit, and the remaining samples, Glucose B and carbon-mixture replicates A and B, were extracted using the mirVana miRNA Isolation Kit (Thermo Fisher Scientific) following the manufacturer’s instructions. For all samples, 23 µL of *Sulfolobus solfataricus* RNA was added as an internal standard during extraction, after addition of the lysis/binding buffer and before subsequent extraction steps, as described previously (Gifford et al., 2016). RNA was eluted in RNase-free water, using 50 µL for the RNeasy extraction and two sequential 50-µL elutions for the mirVana extractions. Initial RNA yield and purity were assessed using a NanoDrop spectrophotometer by measuring absorbance at 260 and 280 nm. Residual DNA was removed with TURBO DNase (Ambion), and final RNA concentrations were quantified using both NanoDrop and a fluorescent RiboGreen assay. Samples were submitted to SeqCenter (Pittsburgh, PA, USA) for library preparation and sequencing. At SeqCenter, samples were treated with Invitrogen DNase to remove residual DNA, and rRNA was depleted using the Ribo-Zero Plus kit. Libraries were prepared using Illumina Stranded Total RNA Prep Ligation with 10-bp unique dual indices and sequenced on an Illumina NovaSeq 6000 platform to generate paired-end 151-bp reads.

### Transcriptome analysis

The forward read alone was used in processing. Raw reads were quality checked using FastQC. Trimmomatic was run under simple mode settings for single end reads on the forward read only, with a sliding window size of 5 bp and required quality score ≥ 20 and reads under 40 bp were discarded. FastQC was run again to identify residual rRNA, a megablast was run against a custom database of rRNA sequences from *Ruegeria pomeroyi* DSS- 3 and *Sulfolobus solfataricus*. rRNA sequences with ≥ 90% identity were counted and removed from downstream analysis. *S. solfataricus* internal standard RNA was identified with a megablast search, and sequences with a ≥ 90% identity were counted and then removed from further analysis. BowTie2 (Langmead and Salzberg 2012) was used to map reads to the Ruegeria Pomeroyi DSS-3 genome (NCBI GCF_000011965.2) using the end-to-end alignment mode.

HTSeq (Anders, Pyl, and Huber 2015) was used to count mapped reads using stranded no settings, minimum alignment quality score of 10, intersection-nonempty mode, set to nonunique-none, and to ignore/discard secondary and supplementary alignments. Differential gene expression was analyzed using the DESeq2 (Love, Huber, and Anders 2014). The transcripts L^-1^ was divided by cells L^-1^ to obtain transcripts cell^-1^. Metabolic pathways and transporters were found through literature and databases such as BioCyc (Karp et al. 2019) and KEGG (Kanehisa 2000).

## RESULTS

### Growth response of DSS-3 to differing carbon substrates

*R. pomeroyi* was grown in batch culture with either 1 mM D-glucose (6 mM C equivalents) or a carbon mixture treatment containing 6 mM total carbon equivalent (TMAO, DMSP, benzoate, succinate, leucine, and glycerol). The two treatments showed similar overall growth yields but differed modestly in growth dynamics (Fig. 1). In the glucose treatment, DSS-3 remained in exponential phase until 71 hr., whereas the mixed-carbon treatment reached maximum cell density earlier, at approximately 56 hr., before both treatments leveled off at similar stationary-phase densities. Growth rates were also slightly higher in the carbon mixture treatment, averaging 4.56 day⁻¹ compared with 3.65 day⁻¹ in the glucose treatment.

**Figure 1.**
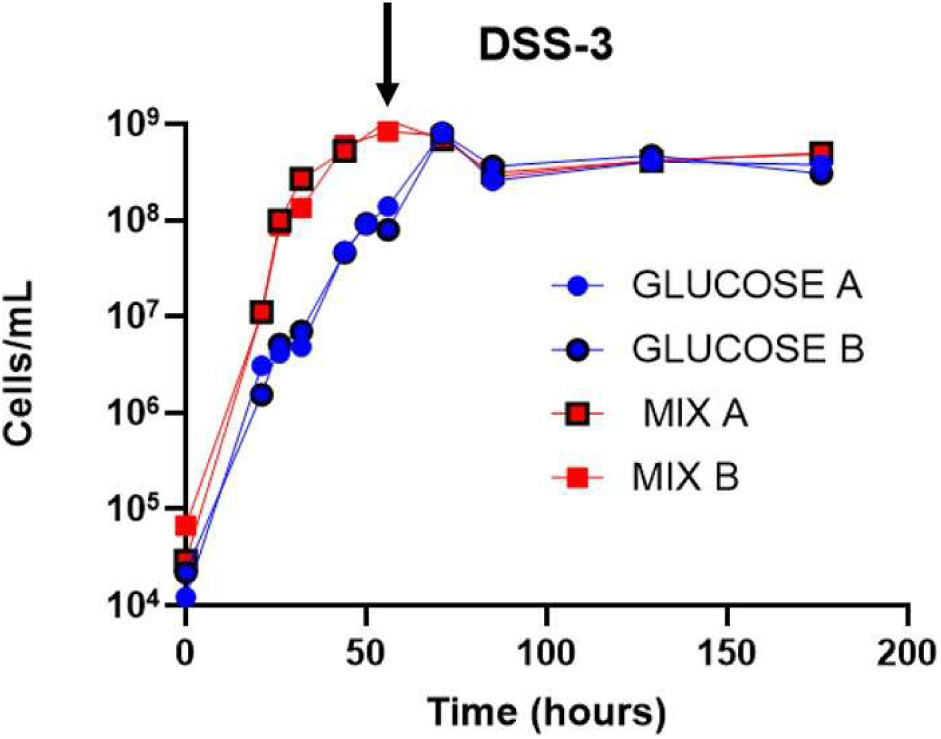
Growth of *Ruegeria pomeroyi* DSS-3 in batch culture with either D-glucose or a mixed-carbon treatment, both at final concentrations of 6 mM total carbon equivalents. The arrow indicates the time point of sampling for RNA collection.

### Genome-wide transcriptional differences

Samples for transcriptome sequencing were collected approximately 50 hr. into the experiment. Significant differential transcription was defined as an adjusted *P* value < 0.05 and an absolute log_2_ fold change > 1. Gene expression differed clearly between the two treatments (Fig. 2). Of the 4,480 genes in *R. pomeroyi*, 339 were significantly enriched in the glucose treatment and 444 were significantly enriched in the carbon-mixture treatment (Table S1). While similar proportions of genes were differentially transcribed in the two treatments, genes enriched in the carbon mixture generally exhibited larger fold changes than those enriched in glucose. The top 50 significantly enriched transcripts for each treatment are shown in Table 1.

**Figure 2.**
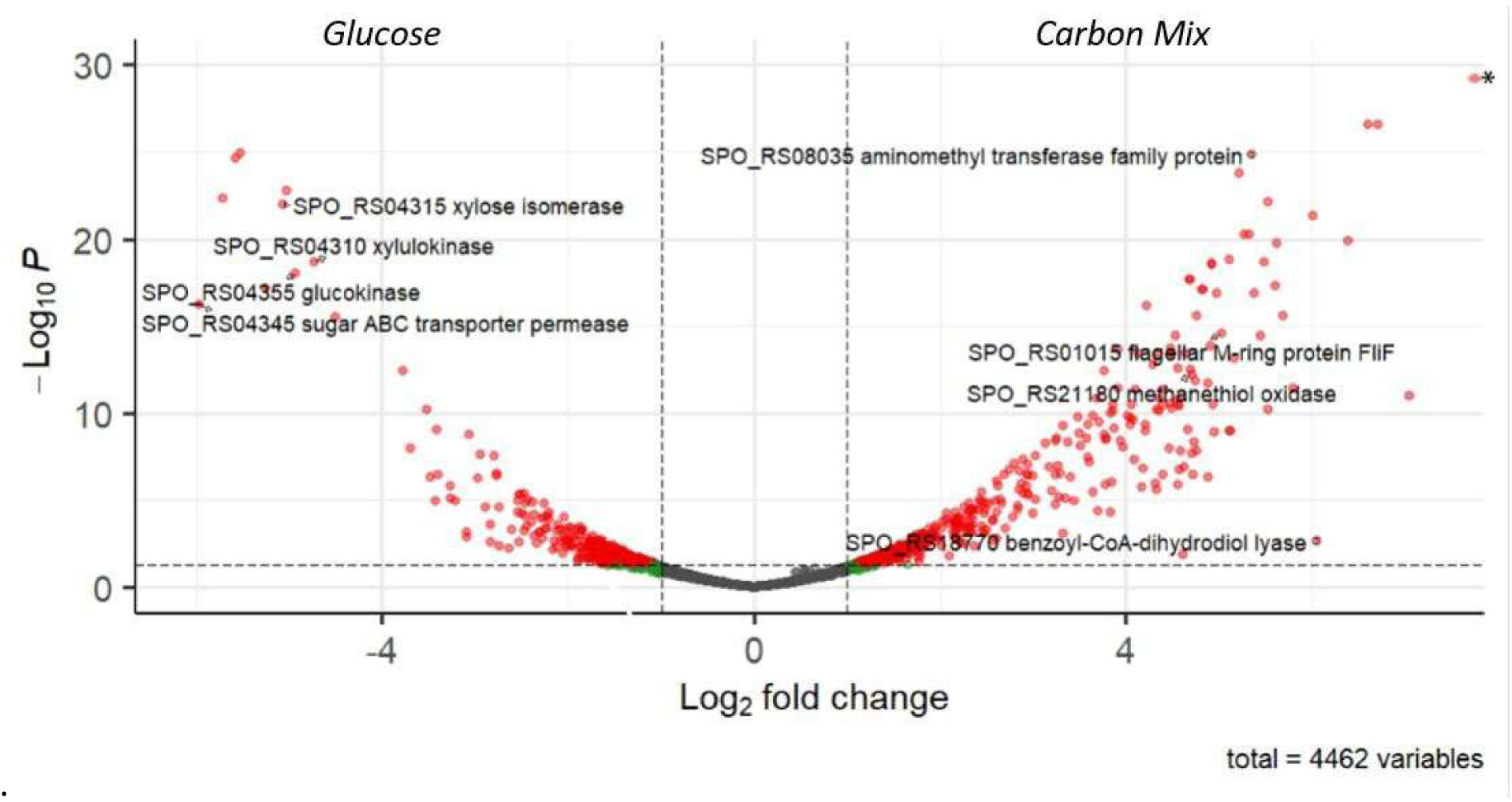
Volcano plot showing differential gene transcription in *Ruegeria pomeroyi* DSS-3 grown in the carbon-mixture treatment relative to the glucose treatment. Positive log2 fold change values indicate genes enriched in the carbon-mixture treatment, and negative values indicate genes enriched in the glucose treatment. The y axis shows statistical significance as −log10 *P*. Red points denote genes with significant differential transcription (adjusted *P* < 0.05 and absolute log2 fold change > 1), green points denote genes with absolute log2 fold change > 1 but adjusted *P* > 0.05, and gray points denote genes with absolute log2 fold change < 1. Selected highly enriched transcripts are labeled. The asterisk indicates a gene with the plotted log2 fold change and an adjusted *P* value of 2.85 × 10⁻³⁹.

**Table 1.** Top 50 enriched transcripts in glucose and carbon mixture treatments.

| Glucose enriched |  |  | C mixture enriched |  |  |
| --- | --- | --- | --- | --- | --- |
| locus tag | gene description | fold change<br>(log2) | locus tag | gene description | fold change<br>(log2) |
| SPO_RS04345 | sugar ABC transporter permease | -6.0 | SPO_RS01665 | EAL domain-containing | 7.1 |
| SPO_RS04340 | D-xylose ABC transporter substrate-binding | -5.7 | SPO_RS22830 | hypothetical | 7.1 |
| SPO_RS04325 | cytochrome-c peroxidase | -5.7 | SPO_RS07910 | ammonium transporter | 6.8 |
| SPO_RS04320 | galactose mutarotase | -5.4 | SPO_RS19245 | thiamine pyrophosphate-dependent dehydrogenase | 6.6 |
| SPO_RS04350 | sugar ABC transporter ATP-binding | -5.2 | SPO_RS18770 | benzoyl-CoA-dihydrodiol lyase | 6.1 |
| SPO_RS04315 | xylose isomerase | -5.1 | SPO_RS19240 | alpha-ketoacid dehydrogenase subunit beta | 6.0 |
| SPO_RS04360 | Gfo/ldh/MocA family oxidoreductase | -5.0 | SPO_RS21840 | hypothetical | 6.0 |
| SPO_RS04355 | glucokinase | -4.9 | SPO_RS11445 | gene transfer agent family | 5.8 |
| SPO_RS04310 | xylulokinase | -4.7 | SPO_RS04680 | response regulator | 5.7 |
| SPO_RS10825 | hypothetical | -4.5 | SPO_RS21175 | IcIR family transcriptional regulator | 5.7 |
| SPO_RS23175 | PaxA | -3.7 | SPO_RS11490 | DNA-packaging | 5.6 |
| SPO_RS05385 | transporter substrate-binding domain-containing | -3.7 | SPO_RS09695 | ester cyclase | 5.5 |
| SPO_RS10815 | porin family | -3.5 | SPO_RS07915 | NAD(P)H-dependent oxidoreductase subunit E | 5.5 |
| SPO_RS06585 | phasin, PhaP | -3.5 | SPO_RS07980 | malate--CoA ligase subunit beta | 5.5 |
| SPO_RS03540 | type I glyceraldehyde-3-phosphate dehydrogenase | -3.4 | SPO_RS18860 | hypothetical | 5.4 |
| SPO_RS01130 | alanine dehydrogenase | -3.4 | SPO_RS21830 | hypothetical | 5.4 |
| SPO_RS07000 | hypothetical | -3.4 | SPO_RS19235 | acetoin dehydrogenase | 5.4 |
| SPO_RS05380 | transporter substrate-binding domain-containing | -3.2 | SPO_RS00360 | ComF family | 5.4 |
| SPO_RS05390 | transporter substrate-binding domain-containing | -3.2 | SPO_RS08035 | aminomethyl transferase family | 5.3 |
| SPO_RS21585 | aldo/keto reductase | -3.1 | SPO_RS07895 | NAD(P)/FAD-dependent oxidoreductase | 5.3 |
| SPO_RS21575 | hypothetical | -3.0 | SPO_RS21845 | hypothetical | 5.2 |
| SPO_RS04330 | CRTAC1 family | -3.0 | SPO_RS00105 | Hpt domain-containing | 5.2 |
| SPO_RS02700 | calcium/sodium antiporter | -3.0 | SPO_RS19250 | NAD(+)/NADH kinase | 5.2 |
| SPO_RS22375 | IS5 family transposase | -3.0 | SPO_RS00275 | DNA repair RadC | 5.2 |
| SPO_RS11605 | porin | -2.9 | SPO_RS11465 | hypothetical | 5.2 |
| SPO_RS05580 | hypothetical | -2.9 | SPO_RS07975 | alanine--glyoxylate aminotransferase | 5.1 |
| SPO_RS05395 | FHA domain-containing | -2.8 | SPO_RS01015 | flagellar M-ring FliF | 5.0 |
| SPO_RS22140 | serine/threonine kinase | -2.8 | SPO_RS01785 | hypothetical | 4.9 |
| SPO_RS05425 | dynamain family | -2.8 | SPO_RS18755 | benzoate-CoA ligase family | 4.9 |
| SPO_RS02545 | DUF1127 domain-containing | -2.7 | SPO_RS19215 | carbohydrate ABC transporter permease | 4.9 |
| SPO_RS22195 | hypothetical | -2.7 | SPO_RS21855 | RebB family R body | 4.9 |
| SPO_RS12770 | hypothetical | -2.7 | SPO_RS05945 | sensor domain-containing diguanylate cyclase | 4.9 |
| SPO_RS01420 | tRNA-Lys-2 | -2.6 | SPO_RS08520 | DUF1153 domain-containing | 4.9 |
| SPO_RS12025 | alanine:cation symporter family | -2.6 | SPO_RS11455 | DUF3168 domain-containing | 4.9 |
| SPO_RS08570 | DUF1127 domain-containing | -2.6 | SPO_RS17520 | flagellin | 4.8 |
| SPO_RS17705 | mechanosensitive ion channel | -2.6 | SPO_RS11495 | hypothetical | 4.8 |
| SPO_RS21590 | carboxymuconolactone decarboxylase family | -2.6 | SPO_RS11425 | DUF2163 domain-containing | 4.8 |
| SPO_RS05420 | dynamain family | -2.5 | SPO_RS07995 | phosphoenolpyruvate carboxylase | 4.8 |
| SPO_RS04335 | ROK family transcriptional regulator | -2.5 | SPO_RS11460 | phage head closure | 4.8 |
| SPO_RS17780 | 50S ribosomal L10 | -2.5 | SPO_RS19230 | carboxymuconolactone decarboxylase family | 4.8 |
| SPO_RS01330 | DMT family transporter | -2.5 | SPO_RS22635 | DUF779 domain-containing | 4.7 |
| SPO_RS03935 | phosphonate ABC transporter ATP-binding | -2.5 | SPO_RS21180 | methanethiol oxidase | 4.7 |
| SPO_RS11575 | 50S ribosomal L9 | -2.5 | SPO_RS01020 | flagellar basal body-associated FliL family | 4.7 |
| SPO_RS12350 | TRAP transporter large permease subunit | -2.5 | SPO_RS11420 | peptidase | 4.7 |
| SPO_RS19400 | 50S ribosomal L23 | -2.4 | SPO_RS07985 | succinate--CoA ligase subunit alpha | 4.7 |
| SPO_RS18275 | DUF1611 domain-containing | -2.4 | SPO_RS07990 | D-glycerate dehydrogenase | 4.6 |
| SPO_RS03535 | LysR family transcriptional regulator | -2.4 | SPO_RS21185 | SCO family | 4.6 |
| SPO_RS15715 | bifunctional methylenetetrahydrofolate dehydrogenase | -2.4 | SPO_RS01005 | FliM/FliN family flagellar motor switch | 4.6 |
| SPO_RS05965 | DUF1127 domain-containing | -2.4 | SPO_RS11435 | phage tail tape measure | 4.6 |
| SPO_RS16600 | orotidine-5'-phosphate decarboxylase | -2.4 | SPO_RS18775 | benzoyl-CoA 2,3-epoxidase subunit BoxB | 4.6 |

### Glucose treatment transcription

To identify the metabolic processes underlying the separation between treatments, we first examined transcripts enriched in the glucose treatment. Overall, genes associated with glucose uptake and catabolism were more highly transcribed in glucose-grown cells than in the carbon-mixture treatment (Fig. 3). In particular, the experimentally verified glucose transporter *xylFGH* was strongly enriched in the glucose treatment, with all three subunits showing transcriptional enrichment relative to the carbon mixture and the substrate-binding subunit *xylF* having the highest relative transcript abundance (Fig. 3B,D).

**Figure 3.**
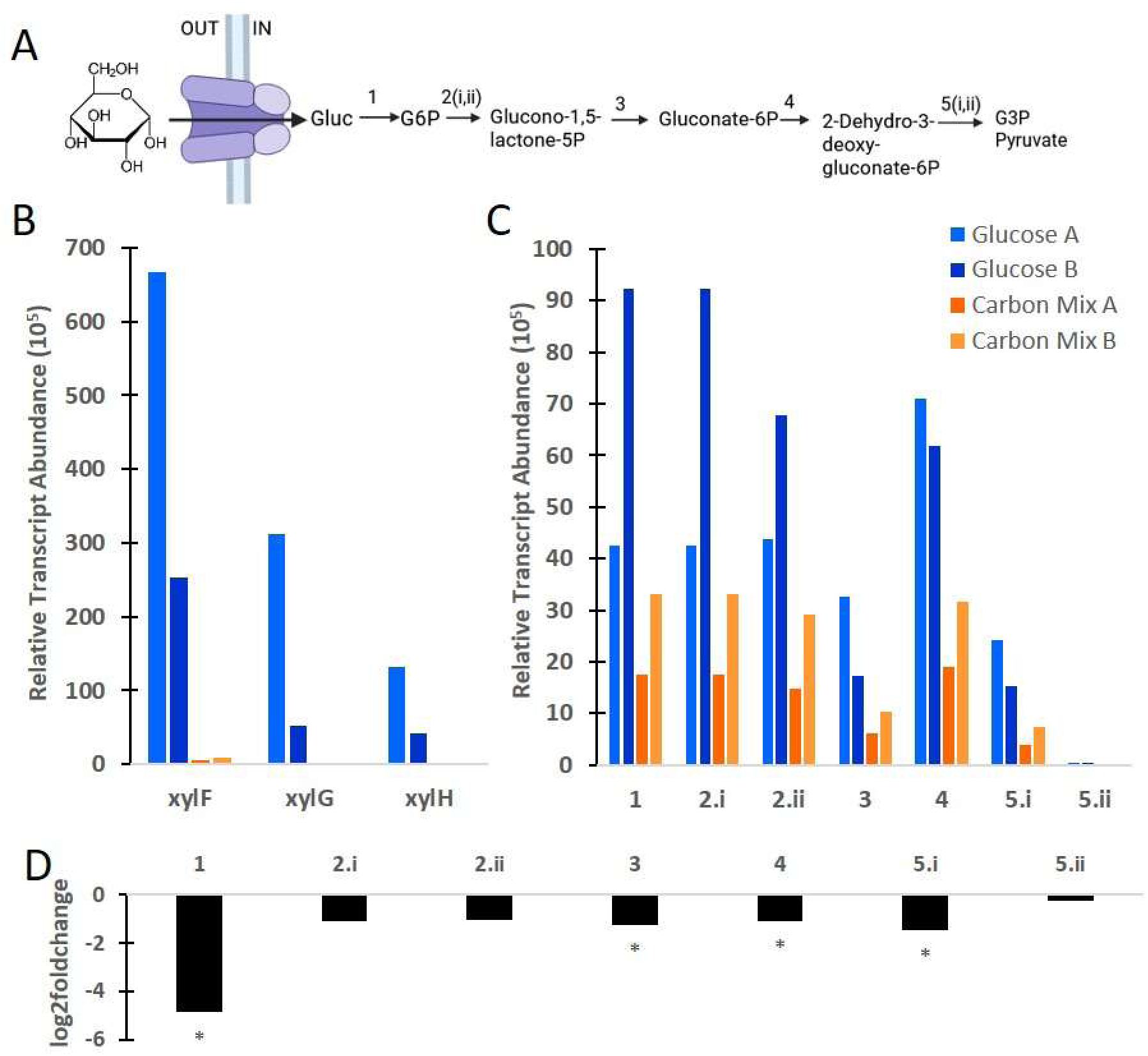
Glucose transport and Entner-Doudoroff pathway transcription in *Ruegeria pomeroyi* DSS-3. **(A)** Schematic of glucose uptake and metabolism through the Entner-Doudoroff pathway. Numbered steps correspond to the genes shown in panels C and D. **(B)** Relative transcript abundance of the glucose transporter (*xylFGH*) components *xylF*, substrate-binding protein; *xylG*, permease; *xylH*, ATP-binding cassette component. **(C)** Relative transcript abundance of each step in the pathway shown in panel A. Steps with multiple enzymes or subunits are shown individually. **(D)** Log2 fold change of transporter components and pathway genes in the carbon-mixture treatment relative to the glucose treatment. Negative values indicate enrichment in glucose treatment, and positive indicate enrichment in carbon-mixture. Asterisks denote genes with adjusted *P* < 0.05. Gene annotations for numbered steps are as follows: 2.i, glucose-6-phosphate dehydrogenase (SPO2048); 2.ii, glucose-6-phosphate dehydrogenase (SPO3033); 5.i, bifunctional 4-hydroxy-2-oxoglutarate aldolase/2-dehydro-3-deoxy-phosphogluconate aldolase (SPO3031); 5.ii, bifunctional 4-hydroxy-2-oxoglutarate aldolase/2-dehydro-3-deoxy-phosphogluconate aldolase (SPOA0330).

Transcriptional enrichment in the glucose treatment was also evident across the early steps of glucose metabolism. Most genes in the Embden-Meyerhof-Parnas pathway trended toward enrichment in glucose-grown cells, but the clearest signal was in the Entner-Doudoroff shunt, for which all genes showed a trend toward higher transcription in the glucose treatment and most were significantly enriched (Fig. 3C,D). The first steps of glucose activation, including glucokinase and glucose-6-phosphate dehydrogenase, showed particularly strong transcript abundances in the glucose treatment. By contrast, enrichment became weaker farther downstream in the pathway, with one of the two aldolase genes showing little differential transcription between treatments (Fig. 3D). These patterns suggest that DSS-3 primarily routed glucose through the Entner-Doudoroff pathway under our experimental conditions rather than maintaining strong transcription across the full glycolytic pathway.

In contrast, genes of the tricarboxylic acid cycle did not show broad enrichment in the glucose treatment. This indicates that the clearest transcriptional distinction in glucose-grown cells was at the level of glucose transport and its initial catabolic steps rather than in downstream central metabolism. In addition to carbon metabolism genes, the glucose treatment was characterized by enrichment of numerous ribosomal RNA transcripts, consistent with the greater overall transcript abundance observed in these samples (Table 1, Table S1). Together, these data show that the glucose treatment was marked by strong transcriptional enrichment of the glucose transporter and early steps of glucose catabolism, particularly the Entner-Doudoroff pathway.

### Transcript enrichment in the carbon mixture

In contrast to the glucose treatment, the mixed-carbon transcriptome was dominated by genes associated with C1 metabolism, likely reflecting the presence of the methylated compounds DMSP and TMAO. Additional enriched transcripts were related to sulfur oxidation, flagellar biosynthesis, leucine biosynthesis, and benzoate oxidation. Many of these genes mapped directly to substrates added in the carbon mixture or to pathways involved in their subsequent metabolism.

#### TMAO

TMAO metabolism showed one of the clearest transcriptional responses in the mixed-carbon treatment. All components of the TMAO pathway were significantly enriched, including all three subunits of the TMAO ABC transporter, *tmoXWV* (Fig. 4). Enrichment extended beyond TMAO transport and initial demethylation to downstream methylated amine oxidation steps. All four *mgd* genes encoding sarcosine oxidase subunits (*mgdA-D*) were enriched in the carbon-mixture treatment (Fig. 4), indicating increased transcription of the pathway linking methylated amine catabolism to C1 metabolism. Because TMAO metabolism generates C1 intermediates through reduction of methylene-tetrahydrofolate, we also observed significant enrichment of genes involved in formate oxidation and C1 assimilation. These included formate dehydrogenase subunits (*fdh*), an *fdhF* formate dehydrogenase subunit, and an associated oxidoreductase. Significantly enriched C1 assimilation genes included *metF*, encoding methylenetetrahydrofolate reductase, and serine hydroxymethyltransferase, which can route C1 units into the serine cycle for biomass production.

**Figure 4.**
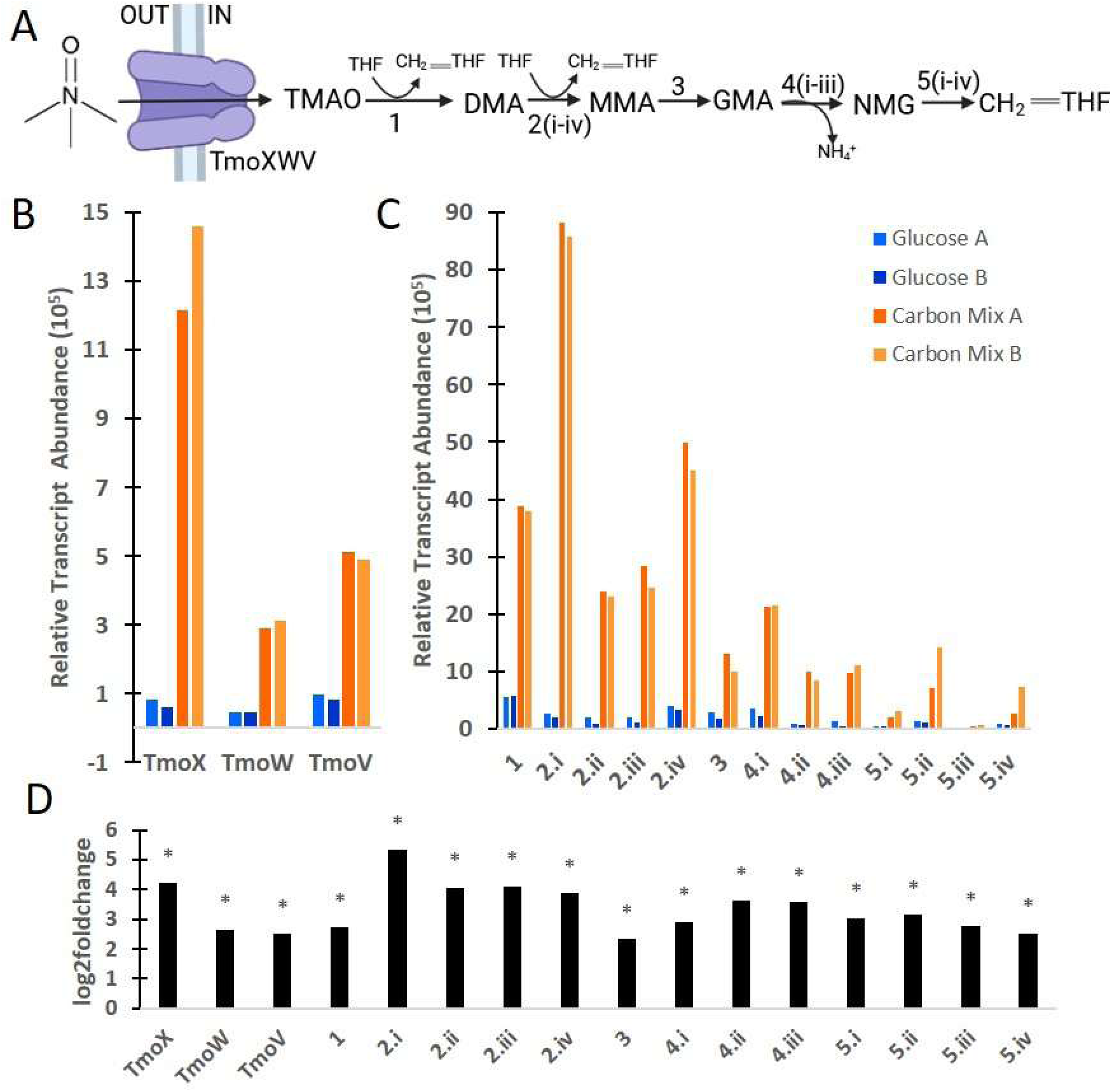
Transcription of TMAO transport and catabolism in *Ruegeria pomeroyi* DSS-3. **(A)** Schematic of TMAO uptake and metabolism. Numbered steps correspond to the genes shown in panels C and D. **(B)** Relative transcript abundance of the TMAO ABC transporter components *tmoXWV*. *tmoX*, substrate-binding protein; *tmoW*, permease; *tmoV*, ATP-binding cassette component. **(C)** Relative transcript abundance of genes associated with each step in the pathway shown in panel A. Steps with multiple enzymes or subunits are shown separately. **(D)** Log2 fold change of transporter components and pathway genes in the carbon-mixture treatment relative to the glucose treatment. Positive values indicate enrichment in the carbon-mixture treatment, and negative values indicate enrichment in the glucose treatment. Asterisks denote genes with adjusted *P* < 0.05. Gene annotations for numbered steps are as follows: 2.i, aminomethyltransferase family protein; 2.ii, *dmmA*; 2.iii, oxidoreductase; 2.iv, *dmmC*; 4.i, FMN-binding glutamate synthase family protein (*mgsA*); 4.ii, *glxC* (*mgsB*); 4.iii, glutamine amidotransferase family protein (*mgsC*); 5.i, sarcosine oxidase subunit gamma (*mgdA*); 5.ii, sarcosine oxidase subunit alpha family protein (*mgdB*); 5.iii, sarcosine oxidase subunit delta (*mgdC*); 5.iv, sarcosine oxidase subunit beta family protein (*mgdD*).

#### DMSP

DMSP metabolism showed a more mixed transcriptional response than TMAO. The two known DMSP transport systems, a BCCT family transporter and the ABC transporter *dmpXWV* (Li et al., 2023; Moran et al., 2012) both trended toward enrichment in the carbon-mixture treatment but were not significantly enriched. Within the ABC transporter, the substrate-binding subunit *dmpX* showed the highest log_2_ fold change and relative transcript abundance in the carbon mixture (Table S1).

Genes from both the DMSP cleavage and demethylation pathways were enriched in the carbon mixture, although the strongest signal was limited to a few key steps (Fig. 5). In the cleavage pathway, most genes trended toward higher transcription in the carbon mixture, but only *dddW*, which catalyzes the first step converting DMSP to acrylate and dimethylsulfide, was significantly enriched. In the demethylation pathway, the only significantly enriched gene was *mtoX*, the terminal step degrading methanethiol to sulfide, formaldehyde, and hydrogen peroxide (Fig. 5). Thus, unlike TMAO, the DMSP signal was not characterized by broad enrichment across the full transport and catabolic pathway, but rather by transcriptional enrichment at specific pathway steps.

**Figure 5.**
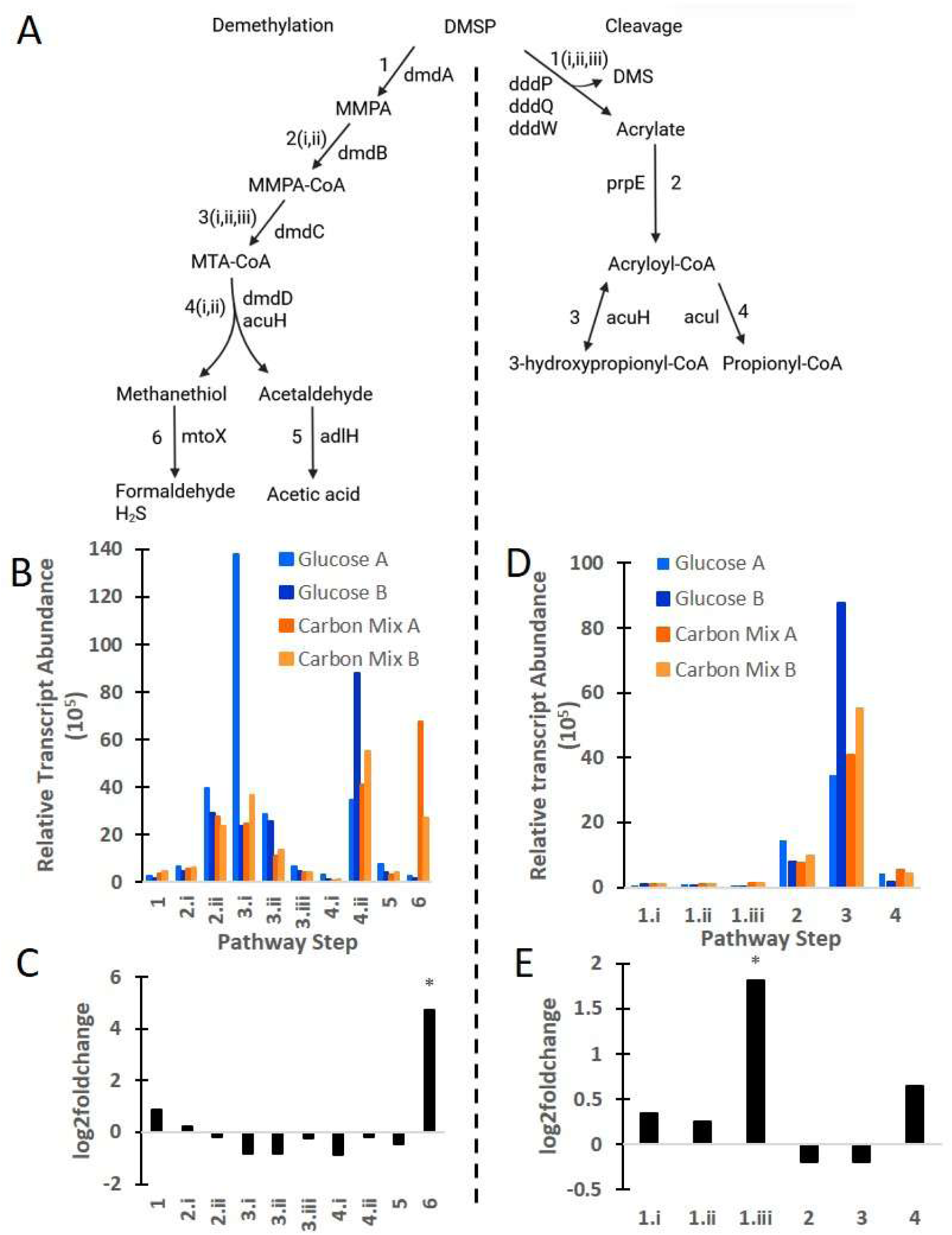
**(A)** DMSP demethylation pathway (left) and DMSP cleavage pathway (right). **(B)** Relative transcript abundance of each metabolic step of the demethylation pathway, with each number corresponding to the metabolic step in panel A. Steps with multiple enzymes/subunits are separated. 2i: acyl-CoA synthetase, 2ii: 3-methylmercaptopropionyl-CoA ligase, 3i: acyl-CoA dehydrogenase C-terminal domain-containing protein, 3ii: acyl-CoA dehydrogenase, 3iii: 3-methylmercaptopropionyl-CoA dehydrogenase, 4i: methylthioacryloyl-CoA hydratase, 4ii: enoyl-CoA hydratase. **(C)** Log2 fold change for each metabolic step in the demethylation pathway. **(D)** Relative transcript abundance of each metabolic step in the cleavage pathway, with each number corresponding to the metabolic step in panel A. Steps with multiple enzymes/subunits are separated. 1i: aminopeptidase P family protein (*ddd*P), 1ii: cupin domain-containing protein (*ddd*Q), 1iii: cupin domain-containing protein (*ddd*W) **(E)** Log2 fold change for each metabolic step in cleavage pathway. Positive log2 fold changes indicate transcript enrichment in the carbon mix, negative indicates enrichment in the glucose treatment. Steps with a padj<0.05 are denoted with an asterisk.

Downstream metabolism of DMSP-derived products was further supported by enrichment of sulfur oxidation genes (Ghosh 2009). These included *soxXVSF* and three cytochrome *c* genes associated with thiosulfate oxidation (Table S1). Additional sulfur oxidation genes, including *soxB* and other cytochrome *c*-related genes, also trended toward enrichment but were not significant. Because DMSP metabolism, like TMAO metabolism, can generate C1 intermediates, some of the enriched downstream C1 oxidation and assimilation transcripts may reflect contributions from both pathways.

### Benzoate

Benzoate oxidation showed a strong transcriptional response in the mixed-carbon treatment, driven by significant enrichment of a core 11-gene operon (Fig. 6). This operon includes two TRAP transporter genes, *boxABC*, *badA-1*, feruloyl-CoA synthase, transcriptional regulators, and a hypothetical protein. All genes within the operon were significantly enriched, including those associated with the first three steps of benzoate oxidation. In contrast, several downstream genes in the *box* pathway located outside this operon, including two enoyl-CoA hydratases and *pcaF*, were not significantly enriched (Fig. 6C). Thus, the benzoate response was concentrated in the operon encoding the early steps of aerobic benzoate degradation rather than distributed uniformly across the full pathway. Relative transcript abundance also differed between the two mixed-carbon replicates, with replicate B showing higher abundance for the first three benzoate oxidation steps than replicate A (Fig. 6B). Although the operon contains two significantly enriched TRAP transporter genes, these genes have not been shown to transport benzoate, and the mechanism of benzoate uptake in DSS-3 remains unresolved.

**Figure 6.**
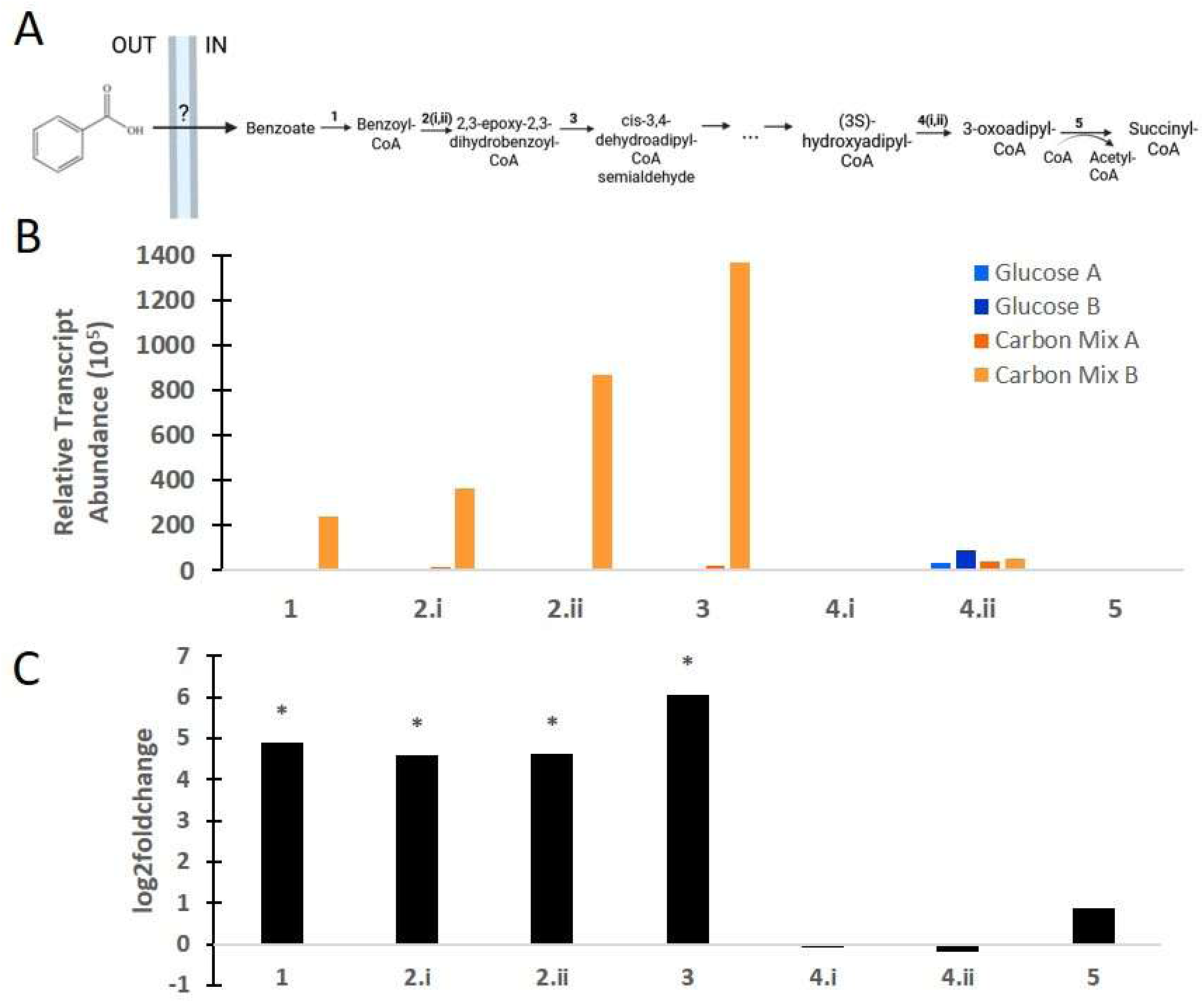
Transcription of aerobic benzoate degradation genes in *Ruegeria pomeroyi* DSS-3. **(A)** Schematic of the aerobic benzoate oxidation pathway. Numbered steps correspond to the genes shown in panels B and C. The transporter responsible for benzoate uptake in DSS-3 is currently unknown. **(B)** Relative transcript abundance of genes associated with each step in the pathway shown in panel A. Steps with multiple enzymes or subunits are shown separately. **(C)** Log2 fold change of pathway genes in the carbon-mixture treatment relative to the glucose treatment. Positive values indicate enrichment in the carbon-mixture treatment, and negative values indicate enrichment in the glucose treatment. Asterisks denote genes with adjusted *P* < 0.05. Gene annotations for numbered steps are as follows: 2.i, benzoyl-coenzyme A 2,3-epoxidase subunit *boxA* (SPO3703); 2.ii, benzoyl-coenzyme A 2,3-epoxidase subunit *boxB* (SPO3701); 4.i, enoyl-CoA hydratase (SPO1687); 4.ii, enoyl-CoA hydratase (SPO0147).

### Succinate and glycerol

In contrast to TMAO, DMSP, and benzoate, succinate and glycerol showed little evidence of a distinct transcriptional response in the carbon-mixture treatment. For succinate, transcription of the experimentally verified TRAP transporter did not differ significantly between treatments, and most genes in the tricarboxylic acid cycle were similarly unchanged (Fig. 7). The only significantly enriched gene associated with succinate metabolism was *sucD*, encoding a succinyl-CoA ligase subunit. Overall, these results indicate that succinate did not produce a strong or coordinated transcriptional signature in the carbon mixture.

**Figure 7.**
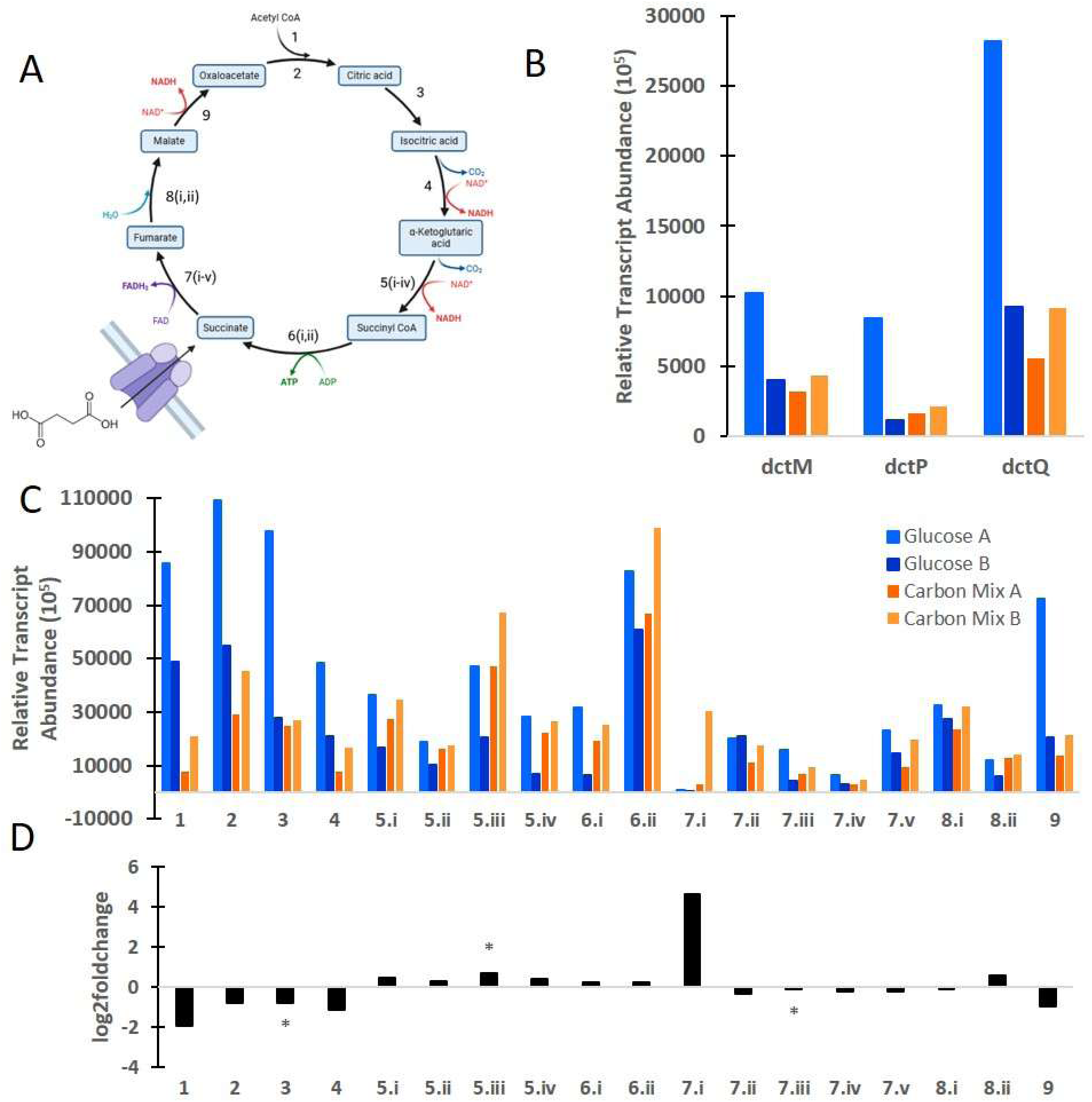
Transcription of succinate transport and tricarboxylic acid cycle genes in *Ruegeria pomeroyi* DSS-3. **(A)** Schematic of the tricarboxylic acid cycle with the succinate TRAP transporter. Numbered steps correspond to the genes shown in panels C and D. **(B)** Relative transcript abundance of the succinate TRAP transporter components. *dctM*, large permease subunit; *dctP*, substrate-binding subunit; *dctQ*, small permease subunit. **(C)** Relative transcript abundance of genes associated with each step in the pathway shown in panel A. Steps with multiple enzymes or subunits are shown separately. **(D)** Log2 fold change of transporter components and pathway genes in the carbon-mixture treatment relative to the glucose treatment. Positive values indicate enrichment in the carbon-mixture treatment, and negative values indicate enrichment in the glucose treatment. Asterisks denote genes with adjusted *P* < 0.05. Gene annotations for numbered steps are as follows: 5.i, dihydrolipoyl dehydrogenase (SPO0340); 5.ii, dihydrolipoyl dehydrogenase (SPO2222); 5.iii, 2-oxoglutarate dehydrogenase E1 component; 5.iv, 2-oxoglutarate dehydrogenase complex dihydrolipoyllysine-residue succinyltransferase; 6.i, succinyl-CoA ligase subunit alpha; 6.ii, ADP-forming succinyl-CoA ligase subunit beta; 7.i, succinate dehydrogenase, cytochrome *b* 556 subunit; 7.ii, succinate dehydrogenase, hydrophobic membrane anchor protein; 7.iii, hypothetical protein; 7.iv, succinate dehydrogenase iron-sulfur subunit; 8.i, fumarate hydratase; 8.ii, class II fumarate hydratase.

Glycerol showed a similarly weak response. Neither the experimentally verified glycerol ABC transporter nor the core glycerol catabolic genes *glpK* and *glpD* were significantly enriched in the carbon-mixture treatment relative to glucose (Fig. 8). Thus, unlike TMAO and benzoate, neither succinate nor glycerol generated a clear transcriptional signal at the level of transport or early catabolism under these conditions.

**Figure 8.**
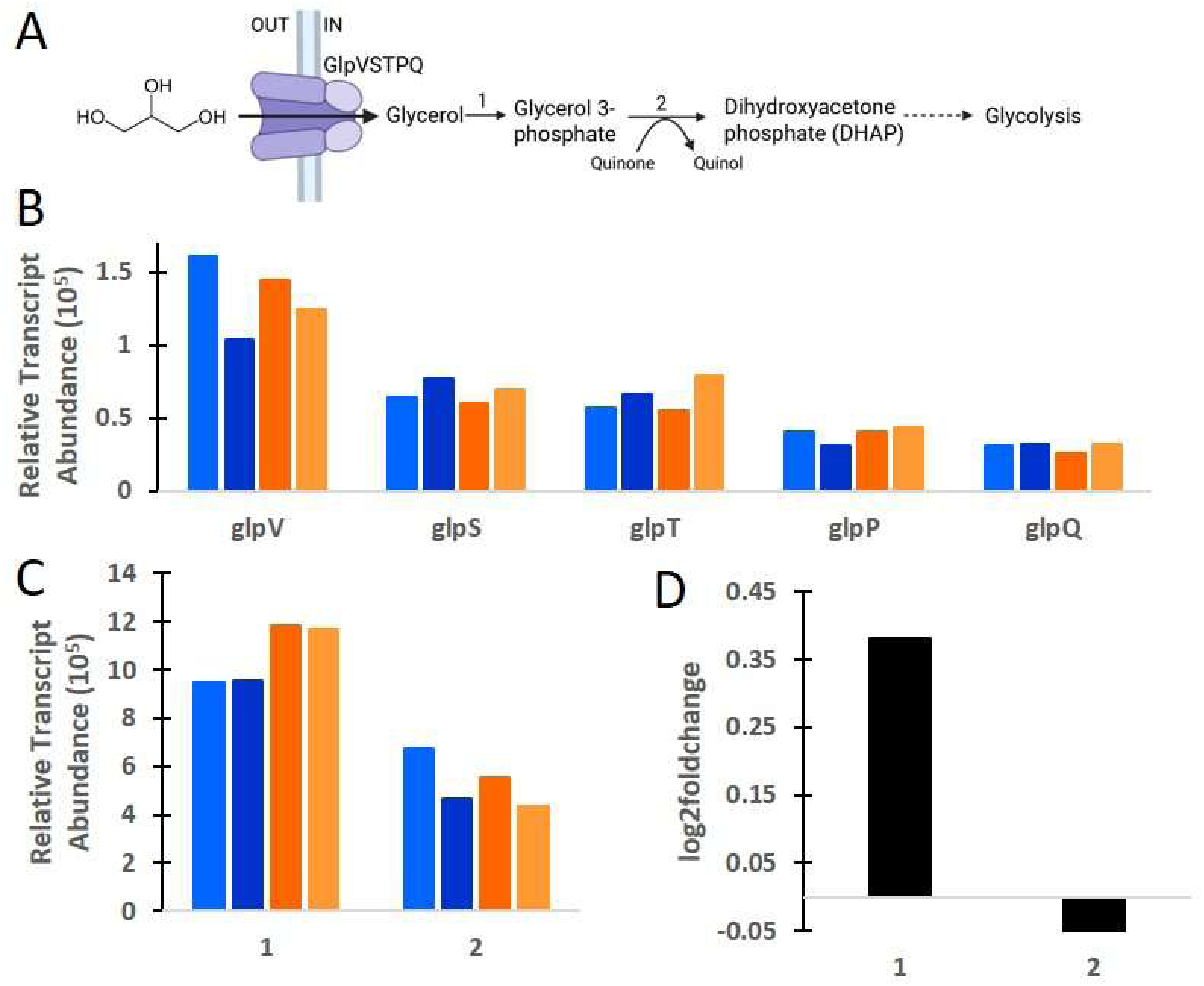
Transcription of glycerol transport and catabolism genes in *Ruegeria pomeroyi* DSS-3. **(A)** Schematic of glycerol uptake and metabolism. Numbered steps correspond to the genes shown in panels C and D. **(B)** Relative transcript abundance of the glycerol transporter components *glpVSTPQ*. **(C)** Relative transcript abundance of genes associated with each step in the pathway shown in panel A: step 1, *glpK*; step 2, *glpD*. **(D)** Log2 fold change of pathway genes in the carbon-mixture treatment relative to the glucose treatment. Positive values indicate enrichment in the carbon-mixture treatment, and negative values indicate enrichment in the glucose treatment. Asterisks denote genes with adjusted *P* < 0.05

### Flagellar assembly

In addition to genes directly associated with substrate transport and catabolism, the mixed-carbon treatment was characterized by strong enrichment of flagellar biosynthesis genes. Of the 30 flagellar genes in DSS-3, 27 were significantly enriched in the carbon mixture treatment (Table S1). Also significantly enriched were homologues of *cckA*, *chpT*, and *ctrA*, regulators best known for roles in cell cycle control but also implicated in flagellar biosynthesis in a related Roseobacter (Zan et al. 2013). These results indicate that the mixed-carbon treatment elicited not only substrate-specific metabolic responses, but also a broader transcriptional response associated with motility.

## DISCUSSION

The genome-wide transcriptional profile of *Ruegeria pomeroyi* DSS-3 differed markedly between a single-substrate environment and a complex carbon mixture. The carbon-mixture treatment was characterized by strong enrichment of C1-related metabolism, consistent with the presence of methylated compounds such as DMSP and TMAO. Overall, these results support the idea that transcriptomics can provide useful insight into substrate availability, as proposed in prior work on marine bacterial metabolism and environmental metatranscriptomics (Poretsky et al. 2010; Gifford et al., 2011; Gifford et al. 2013; Moran et al. 2013; Nowinski and Moran 2021). However, the strength and specificity of this signal depended strongly on pathway position and metabolic context. In general, transcriptional enrichment was strongest at the first committed steps of substrate utilization and weakened as metabolism converged on central pathways. In addition, the presence of multiple substrates appeared to blur transcriptional signatures that are clearer in simpler substrate systems, emphasizing the challenge of applying transcript-based inference to chemically complex DOC pools in natural environments.

Before evaluating which genes provided the clearest substrate-linked signals, it is important to note that the two treatments were sampled at the same time but likely not at the exact same physiological state: the carbon-mixture cultures had reached stationary phase, whereas the glucose cultures were still in late exponential phase. Because bacterial transcriptional profiles shift substantially across growth phases (Deng et al. 2021; Veselovsky et al. 2022; Lim et al. 2025), some observed differences likely reflect physiology in addition to substrate-specific metabolism. This is consistent with the higher transcript inventories in the glucose treatment and the enrichment of ribosomal RNA and ATP synthase-related transcripts, both associated with exponential growth. It is therefore possible that some metabolic responses in the carbon-mixture treatment were underrepresented relative to what would have been observed earlier in growth. Nevertheless, the recovery of coherent transcriptional signatures for glucose, TMAO, benzoate, sulfur oxidation, and C1 metabolism indicates that the major substrate-linked patterns were not driven solely by growth phase but reflected meaningful responses to the supplied carbon substrates.

### Early pathway genes were the best indicators of substrate availability

Across multiple substrates, the clearest transcriptional signals occurred at pathway entry points, where enzymes are most specific to the supplied compound and, therefore, most directly reflect its uptake and initial processing. This pattern was evident for glucose, TMAO, and benzoate. In the glucose treatment, enrichment was strongest in the early steps of glycolysis and the Entner-Doudoroff pathway, whereas in the carbon mixture, the TMAO transporter and initial catabolic genes were strongly enriched. For benzoate, *badA*-1 and *box*ABC were among the clearest indicators of substrate use. These results indicate that the most reliable transcriptomic markers of substrate availability are genes involved in the first committed transformations of a substrate rather than genes acting later in its assimilation. This interpretation is consistent with previous observations that early pathway genes can provide robust signals of substrate use in both isolates and environmental datasets. For example, *tdm*, encoding the first step of TMAO metabolism, is highly expressed in marine bacteria and ocean metatranscriptomes (Lidbury et al., 2014), and *box*A and *box*B have been identified as key markers of benzoate degradation in oil-exposed marine communities (Knapik et al. 2020).

A likely explanation is that metabolic connectivity increases farther downstream in a pathway. Entry-point enzymes typically interact with a narrow range of substrates and therefore retain greater substrate specificity in their transcriptional response. By contrast, downstream enzymes are more often connected to central metabolism and may process intermediates shared across multiple pathways, reducing their value as indicators of any one substrate. The benzoate pathway illustrates this pattern well: the initial *box* genes were strongly enriched in the carbon mixture, whereas later steps such as enoyl-CoA hydratases, *pca*F, and *suc*D showed weaker or nonsignificant responses. Similar logic applies to DMSP metabolism, where transcriptional enrichment occurred at the beginning and end of the pathway but was weaker across many intermediate steps. In that case, downstream genes likely reflect shared sulfur- and C1-processing steps that can also be fed by other substrates in the mixture, particularly TMAO. Thus, while downstream genes may still indicate broader metabolic themes such as sulfur oxidation or C1 assimilation, they are less effective for resolving which specific substrate is driving the response.

### Transporter transcription is a complex signal

Because transporters mediate the first direct interaction between the cell and an extracellular substrate, we expected them, particularly substrate-binding components, to be among the strongest transcriptional indicators of substrate availability. This expectation was motivated by previous work suggesting that transporter expression can provide insights into DOC composition and substrate uptake (Poretsky et al. 2010; Schroer et al. 2023). However, this pattern was only partly supported here. Of the experimentally-verified transporters examined, only a subset showed clear enrichment in the treatment containing their substrate. The clearest examples were the glucose transporter in the glucose treatment and the TMAO ABC transporter in the carbon mixture, both of which showed strong enrichment, with the substrate-binding subunit as the most highly transcribed component. In contrast, transporters expected to respond to glycerol, DMSP, leucine, and succinate showed weak or no enrichment. Thus, although transporter transcription sometimes reflected substrate availability, it was not as consistently diagnostic as predicted.

One pattern that was consistent across transporter systems was that the substrate-binding subunit tended to show the strongest response relative to membrane and ATPase or permease components. This was true not only for transporters with strong enrichment, but also for weakly responding systems such as the glycerol and DMSP ABC transporters. Even so, the inconsistent significance of entire transporter systems indicates that direct substrate interaction alone is not sufficient to guarantee a strong transcriptional signal.

Transporter transcription is likely shaped by regulatory processes that decouple transcript abundance from simple substrate presence, including extracellular and intracellular substrate concentrations, constitutive expression, and broader metabolic state. This view is consistent with work showing that bacterial transporters can function within complex signaling and regulatory networks rather than as simple on-off substrate sensors (Piepenbreier, Fritz, and Gebhard 2017; Västermark and Saier 2014). In addition, interpretation depends heavily on accurate functional annotation. In this study we examined 6 of the 18 transporters in DSS-3 experimentally verified by Schroer et al (Schroer et al. 2023), including transporters for glucose, TMAO, glycerol, and DMSP, a succinate TRAP transporter, and a DMSP BCCT family transporter. Many others are annotated primarily through sequence homology, which introduces uncertainty in linking transcriptional enrichment to a specific substrate (Schnoes et al. 2009; Schroer et al. 2023). These factors complicate the use of transporters as straightforward transcriptional indicators of substrate availability and likely help explain why several transporters in our dataset did not conform to our initial expectations.

### Substrate mixtures can blur transcriptional signals seen in single-substrate studies

Comparison with previous single-substrate studies indicates that the presence of multiple carbon compounds can weaken or obscure transcriptional signals that are clearer when DSS-3 is grown on an individual substrate. Earlier work with DSS-3 in single-substrate or substrate-limited systems identified distinct transcriptional responses to compounds such as benzoate, glycerol, succinate, and DMSP (Boulton 2021; Reisch et al. 2013; Table 2). In our carbon-mixture treatment, some previously described signals were retained, whereas others were diminished or lost (Table 2). The clearest example of signal retention was benzoate, for which *badA*-1 and *box*ABC remained strongly enriched, indicating that the first committed steps of benzoate oxidation are robust indicators even in the presence of other carbon sources. In contrast, the glycerol response was much less apparent: genes previously reported to respond to glycerol availability were not significantly enriched despite glycerol being present. Succinate showed a similarly weak response, with little evidence that either its transporter or most TCA cycle genes provided a specific signal of substrate availability. These comparisons suggest that pathway-specific entry-point genes may remain detectable for some substrates in mixed carbon pools, whereas others may become effectively masked.

**Table 2.**
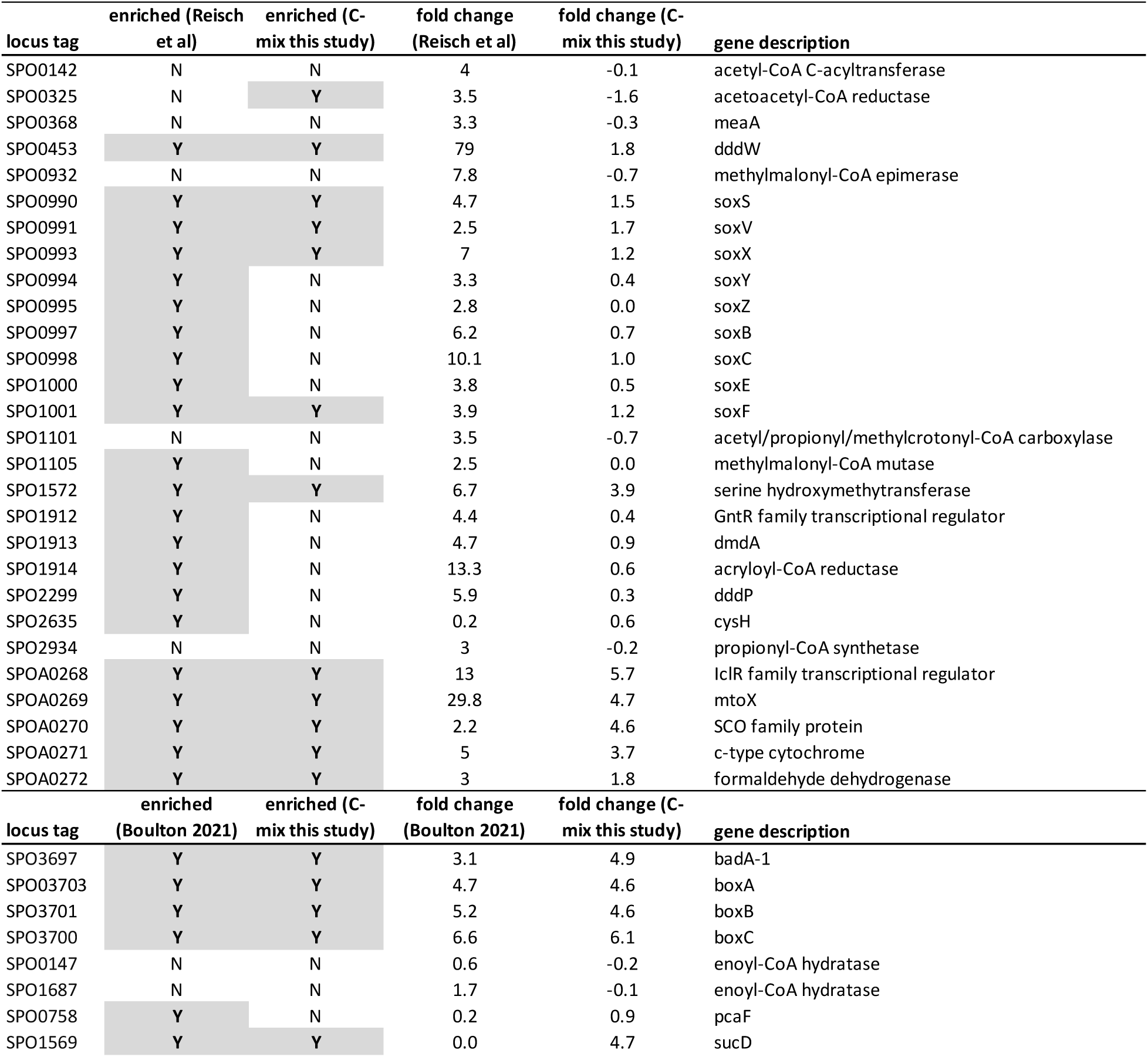
Comparison of transcriptomic response of DSS-3 in the carbon mixture versus Reisch et al. 2013 and Boulton 2021.

DMSP provided a more nuanced case. Some signals observed here were similar to those reported previously, particularly genes involved in sulfur oxidation, formaldehyde oxidation, and C1 assimilation, whereas others were reduced in strength or no longer significant (Table 2). This suggests that a mixed-substrate environment does not eliminate DMSP-associated transcriptional responses but can dilute them and shift interpretation toward broader metabolic themes such as methylated compound and reduced sulfur metabolism rather than substrate-specific inference. This conclusion is also consistent with the fact that DMSP and TMAO metabolism both converge on shared C1-processing pathways.

More broadly, these comparisons support the view that substrate diversity reduces both the magnitude and specificity of transcriptomic signatures. In single-substrate systems, transcriptional responses can be more cleanly linked to one available compound, whereas in mixed-substrate systems the cell integrates multiple substrate inputs simultaneously, producing a more composite metabolic signal. This does not mean transcriptomics loses its value in complex carbon pools, but it does mean that interpretation should focus on the most substrate-specific features of a pathway and avoid over-interpreting weaker downstream responses.

### Indirect transcriptional responses

In addition to substrate-linked signals, the carbon-mixture treatment produced responses that were more difficult to interpret, highlighting that transcriptomes capture broader physiological and ecological responses as well as direct substrate use. One of the clearest examples was the strong enrichment of flagellar biosynthesis genes. DSS-3 showed significant enrichment of 27 of 30 flagellar genes, along with enrichment of *cck*A, *chp*T, and *ctr*A homologues associated in related roseobacters with flagellar biogenesis and motility regulation (Zan et al. 2013).

Because the mixture included compounds such as DMSP, TMAO, and leucine that are commonly associated with phytoplankton exometabolites, one possibility is that DSS-3 responded not only metabolically but behaviorally, interpreting these substrates as cues of a resource-rich microscale environment. This idea is consistent with the broader ecology of roseobacters, which are often associated with phytoplankton and are known to respond to compounds in the phycosphere (González et al. 2000; Miller and Belas 2006; Smriga et al. 2016). Phytoplankton are also known sources of compounds such as DMSP, TMAO, and amino acids (Li et al. 2015; Michael et al. 2016; Krayushkina et al. 2019), and DSS-3-like roseobacters often increase during phytoplankton blooms (Kaur et al. 2018). Thus, some transcriptional responses in the carbon mixture may reflect ecological strategy in addition to catabolism.

### Transcriptional false positives

We also observed significant enrichment of genes whose annotated functions did not map directly onto the supplied compounds. These included transporter genes annotated for glycine betaine/proline, spermidine/putrescine, tungsten, taurine, and oligosaccharides. Such apparent false positives illustrate an important limitation of transcript-based inference. In some cases, these enrichments may reflect inaccurate functional annotation, particularly for transporters assigned on the basis of sequence homology rather than experimental validation (Schnoes et al. 2009; Schroer et al. 2023). In other cases, the annotated transporter may have broader substrate specificity than currently recognized or may be co-regulated with pathways responding to other substrates that were present. For this reason, enrichment of an annotated transporter should not automatically be taken as evidence that its named substrate was available. Instead, these genes are best interpreted cautiously and in conjunction with nearby metabolic genes, operon structure, and experimentally validated function where possible.

Unexpected enrichments were not limited to transporters. We also observed enrichment of pyruvate metabolism genes even though pyruvate was not added to the media. Given pyruvate’s central position in metabolism, these signals may reflect secondary metabolic adjustments, shifts in intracellular metabolite pools, or currently unrecognized pathway linkages rather than direct evidence for extracellular pyruvate availability. More broadly, these results emphasize that transcriptomic responses in mixed-substrate environments contain both direct and indirect information. Transcriptomes should therefore be interpreted as contextual readouts of metabolic state rather than simple inventories of available compounds.

### Conclusion

Overall, our results show that transcriptomics can provide meaningful insight into substrate availability, but the strength and specificity of that signal depend strongly on where a gene sits within a metabolic pathway and on the complexity of the surrounding carbon pool. The most reliable indicators were generally transporters or early pathway genes associated with the first committed steps of substrate use, whereas downstream genes were more difficult to interpret because of their greater metabolic connectivity and overlap with central metabolism. At the same time, the mixed-carbon treatment demonstrated that the presence of multiple substrates can blur transcriptional signals that appear clearer in single-substrate systems, while also generating indirect physiological responses that complicate simple substrate-to-gene inference. These findings suggest that transcriptomic approaches will be most powerful when applied cautiously, with emphasis on experimentally validated transporters and pathway entry-point genes, and when interpreted within the broader context of cellular physiology and environmental complexity.

## ACKNOWLEDGEMENTS

This work was financially supported by the United States National Science Foundation Division of Ocean Sciences (Grant numbers OCE-1850692 and OCE-2505930) to S.M.G. We thank Garrett Sharpe for coding assistance.

